# Evidence for an indigenous female mouse urobiome

**DOI:** 10.1101/2025.08.20.671418

**Authors:** Sidra Sohail, Daniel Bushnell, Mark Khemmani, Sridhar Narla, Olivia Lamana, Bhanu Sharma, Robert B. Moreland, Alan J. Wolfe, Catherine S. Forster

## Abstract

Mice have been used as a valuable model of understanding pathophysiological mechanisms of urinary tract infection for almost six decades. Mice offer many advantages including genetic manipulation to test the role of genes and mechanisms, the availability of germ-free mice, and similarities to humans in innate immune defenses and the strain-dependent presence of vesicoureteral reflux. However, like with humans, the mouse bladder urine above the urinary sphincter has generally been assumed to be sterile. Yet, given the presence of urobiomes in other mammals and the emerging role of the human urobiome in the defense of the urinary bladder and upper urinary tract, the existence of a mouse urobiome should be critically examined as indigenous microbiota may influence experimental results. To determine if an indigenous murine urobiome exists, we obtained expressed urine from two sets of female C57BL/6J mice during three different intervals using two different extraction and sequencing methods and analyzed them simultaneously by a single method. For one set, we also obtained urine by suprapubic aspiration, which we compared to the paired expressed urine samples. We conclude that an indigenous murine urobiome exists and that expressed urine contains post-urethral microbes.

## Introduction

Urinary tract infections (UTI) are among the most common infections in clinical practice (1). Commonly diagnosed with urine dipstick and in office urinalysis, UTIs are responsible for the bulk of antibiotic prescriptions worldwide, leading to antibiotic resistant microbes (2). UTIs, especially when they recur, are associated with renal scarring and can lead to eventual renal failure (3, 4). In one autopsy study of all-cause mortality, healed pyelonephritis was found in 18% of patients while a total of 33% demonstrated evidence of active or healed pyelonephritis (3). Consequently, understanding the pathophysiological mechanisms involved in UTIs is of key importance (5).

Urine has long been believed to be sterile above the urethral sphincter (6, 7). This belief is reinforced by false negative urine dipstick analyses, and the fact that most standard urine cultures (SUC) exhibit no growth in the absence of symptoms. However, much of the myth of urine sterility is due to SUC methods that favor facultative anaerobes, such as members of the genera *Escherichia*, *Pseudomonas*, *Klebsiella*, *Proteus*, *Staphylococcus*, and *Enterococcus*, with *Escherichia coli* by far considered the predominant cause of UTI (5, 6, 8). We now know that human bladder urine is not sterile and that appropriate culture conditions can culture 70% of the genera detected by DNA sequencing methods (8, 9). Furthermore, negative SUC or urine dipstick tests can miss many important emerging uropathogens, as well as lead to unnecessary antibiotic use that destroys the indigenous healthy urinary microbiome (also known as the urobiome) (1).

Research aimed at bettering our understanding of the virulence mechanisms and host-pathogen interactions in UTI have relied on animal models (10). Key in experiments with dogs, cats, rats and mice has been the assumption that urine above the urethral sphincter is sterile and that introduction of test microbes reflects human UTI. Recently, this notion has been disproved with low biomass, indigenous microbiota in dogs (11-14), cats (15) and rats (16, 17). While defining low biomass microbiota brings challenges associated with ruling out contamination (15, 18, 19), the indigenous microbiota may play a role in the defense of the bladder and understanding normal states and dysbiosis are key in any model of UTI, as well as in human disease (20-23).

Mice have been used as a valuable model of understanding pathophysiological mechanisms of UTI for almost six decades (10). Mice offer many advantages including genetic manipulation to test the role of genes and mechanisms, the availability of germ-free mice (which lack microbiota), and similarities to humans in innate immune defenses and the strain-dependent presence of vesicoureteral reflux (10, 24). However, like with humans, the mouse bladder urine above the urinary sphincter has generally been assumed to be sterile. Yet, given the presence of urobiomes in other mammals and the emerging role of the human urobiome in the defense of the urinary bladder and upper urinary tract, the existence of a mouse urobiome should be critically examined as indigenous microbiota may influence experimental results. Indeed, two studies provide some evidence that a mouse urobiome does exist.

One study examined microbes from mouse bladder urine sampled by suprapubic aspirates using aerobic culture methods and viability assays (25). By aerobic culture, they found 150 ± 220 CFU/ml (n=5); however, when cell viability was examined using a BacLight LIVE/DEAD bacterial viability assay, they found 4.5 ± 2.8 x 10^6^ cells/mL. These were scored as “viable but not culturable.” Another study examined the effect of microbiota on the transcriptome and weight of the urinary bladder by comparing germ-free (GF) and specific pathogen-free (SPF)-housed mice (26). The GF urinary bladders had on average 25% lower weight than bladders from mice housed under standard SPF conditions. These experiments were carried out in two different facilities using two different C57BL/6 substrains. Histologically, hematoxylin-eosin-stained sections of GF and SPF bladder tissue showed regular bladder tissue composition. The difference in bladder weight was not correlated to any tissue structural changes by histological assessment of stained sections. To confirm that reduced bladder weight was not a microbiota-driven developmental anomaly, the investigators depleted microbiota among SPF-housed mice using broad-spectrum antibiotics. This continuous antibiotic treatment reproduced the reduced bladder weight phenotype in SPF mice. Thus, reduction in bladder weight was consistent with an adaptive bladder change in response to the absence of microbiota or their products. This study suggests that microbiota can have a profound effect on the host it inhabits. Since germ-free mice are devoid of gut and bladder microbiota, it is impossible to distinguish the effect of urinary microbiota from gut microbiota. However, removing all microbiota from mice affects urinary bladder weight and size. Taken together, these two studies provide a compelling reason to ask whether an indigenous murine urobiome exists.

To answer this question, we obtained expressed urine from two sets of female C57BL/6J mice during three different intervals using two different extraction and sequencing methods and analyzed simultaneously by a single method. For one set, we also obtained urine by suprapubic aspiration, which we compared to the paired expressed urine samples. We conclude that an indigenous murine urobiome exists and that expressed urine contains post-urethral microbes.

## Materials and methods

### Ethics Statement

This study was carried out in strict accordance with the recommendations in the Guide for the Care and Use of Laboratory Animals of the National Institutes of Health. This study was approved by the institutional animal care and use committees at the University of Pittsburgh (protocol 23124147) and the Biomedical Research Institute (protocol 20-01) prior to initiation of the work. Suprapubic aspirations were performed under inhaled isofluorane anesthesia, and all efforts were made to minimize suffering.

#### Mice

Two sets of female C57BL/6J mice (Jackson Labs, Bar Harbor, ME; stock no. 000664) were analyzed. We allowed a three-week acclimation period prior to urine collection for animals in set 1; the mice were 10 weeks at the time of urine collection. For set 2, which were housed in a different institution, we allowed a one-week acclimation period prior to urine collection; the mice were 10 weeks at the time of urine collection. The mice in set 3 were some of the same mice in set 1, but they were aged for an additional 4 months. There were no interventions or other experimental procedures performed in the mice in set 3 while they aged. The mice in set 3 were 27 weeks at the time of collection.

##### Housing and husbandry

All mice were socially housed with 5 mice per cage, with standard enrichment in accordance with guidelines of the Association for Assessment and Accreditation of Laboratory Animal Care. All animals within each set were housed within the same room within the same vivarium and were given the same chow and water from the same source. All mice for this work were commercially-purchased: no mice were bred for this work.

##### Animal Care and Monitoring

Bladder expression for urine is not generally a painful procedure and routine analgesia was not provided. Mice was anesthetized with isoflurane prior to suprapubic aspiration. We had both ketoprofen and buprenorphine available should mice display any signs of pain or distress, but no mice required analgesia. There were no expected or unexpected adverse events. As this was an observational study of the mouse urobiome, the endpoint was obtaining sufficient urine for urobiome analysis. No mice were euthanized as part of this work.

### Urine Collection

#### Expressed Urine

To collect expressed urine samples, we first scruffed each mouse while holding the tail up in the palm of the collector’s hand and used alcohol wipes to clean the perineum. The mouse was then held over the open tube (sterile, DNase and RNase-free microcentrifuge tubes), and suprapubic pressure applied to induce voiding directly into the tube without any contact with the lid or rim. We had to combine urine samples from multiple expressions to obtain sufficient urine volume for sequencing, which we determined was a minimum of 0.5 mL. On average, each voiding produced approximately 50 μL of urine. We added AssayAssure® (nucleic acid preservative, Sierra Molecular Corp, Princeton, NJ, LOT G3030C) to the urine in a 1:9 ratio and froze the sample at -80°C. For sets 1 and 2, only expressed urine was obtained.

#### Suprapubic Aspiration

For set 3, we obtained paired urine samples collected by both expression (as described above) and suprapubic aspiration. To minimize the time between the expressed and aspirated samples for each mouse, we collected the expressed samples immediately before collecting the aspirated samples. For suprapubic aspiration, we anesthetized the mice with inhaled isoflurane prior to aspiration. Once anesthesia was confirmed, we placed the mice on their backs and used Nair^TM^ (Church and Dwight Co, Ewing, NJ) to remove the fur on the suprapubic region. We then thoroughly cleaned the area with alcohol swabs. We placed the ultrasound probe (VisualSonics RMV 710B Ultrasound Imaging Scanhead 25 MHz 70 x 140 μm for Vevo 770, FUJIFILM VisualSonics Inc, Toronto, Canada) on the abdomen with the ultrasound machine (Vevo 770 High-Resolution In Vivo Micro-Imaging System, FUJIFILM VisualSonics Inc, Toronto, Canada) on the maximum refresh rate. Once the bladder was in view, we inserted a 1mL syringe with a 30G needle into the abdomen and into the bladder under ultrasound guidance. From the bladder, we aspirated the urine and transferred it into a sterile Eppendorf tube (Fisher brand Premium 1.5 mL MCT Graduated Natural Eppendorf Tubes, sterile, RNase/DNase free, Cat No. 05-408-129). We added AssayAssure® (Sierra Molecular Corp, Princeton, NJ, LOT G3030C) in a 1:9 ratio and stored the samples at -80°C.

#### Genome Extraction

For sets 1 and 3, each mouse’s multiple samples of urine were thawed at 4°C and then pooled into a single microcentrifuge tube. Next, 0.5 mL of urine was transferred to a deepwell plate, which was centrifuged for 10 mins at 1300 x g and all but 100 μL of supernatant was removed. Enzymatic treatment consisting of 4 mg Lysozyme, 3 kU Mutanolysin, 4 μg Lysostaphin, and 200U Achromopeptidase in buffer (20 mM Tris, 2 mM EDTA, 1.2% Triton, pH 8.0) were added to each sample and incubated at 37°C, 5 hours, shaking intermittently at 1000rpm (Eppendorf Thermomixer with SmartBlock DWP1000). Samples were then processed using the Dneasy 96 Blood & Tissue kit (Qiagen, Germantown, MD) according to the manufacturer’s instructions. To assess potential DNA contamination, extraction negative controls (no urine) were processed with samples and sequenced in every run. For set 2, genomic DNA was isolated using the DNeasy PowerSoil Kit (Qiagen, Germantown, MD), according to kit instructions, with the exception that DNA was eluted from the column with nuclease-free water (Sigma-Aldrich, St. Louis, MO).

#### Sequencing

The urine samples from sets 1 and 3 were sequenced by SeqCenter (Pittsburgh, PA), where they were prepared using Zymo Research’s Quick-16S kit with phased primers targeting the V4 regions of the 16S rRNA gene (5’-GTGYCAGCMGCCGCGGTAA-3’, 5’-GGACTACNVGGGTWTCTAAT-3’) and sequenced. Following clean-up and normalization, samples were sequenced on a P1-600cyc NextSeq2000 Flowcell to generate 2 x 301 bp PE reads. Adapters were trimmed.

The urine samples from set 2 were sequenced by the Genomics and Epigenomic Shared Resource at the Lombardi Comprehensive Cancer Center at Georgetown University. Library preparation and sequencing was performed according to Illumina 16S Metagenomics Sequencing Library Sample Preparation Guide. In brief, genomic DNA isolated from mouse urine samples was used to amplify the 16S V3-V4 region. The primers used for the amplification were 16S Amplicon PCR Forward Primer (5’-TCGTCGGCAGCGTCAGATGTGTATAAGAGACAGCCTACGGGNGGCWGC AG) and 16S Amplicon PCR Reverse Primer (5’-GTCTCGTGGGCTCGGAGATGTGTATAAGAGACAGGACTACHVGGGTATC TAATCC). The IDT-Illumina DNA-RNA UD Indices SetA Tagmentation index kit was used for the index PCR. The amplicons were pooled and sequenced with MiSeq v3 600 cycle reagent kit (Illumina) with 15% PhiX spike-in.

#### Taxonomic Identification

For sets 1, 2, and 3, high-quality reads were obtained by trimming low-quality sequences using Cutadapt (27). The DADA2 software (28) was then used to process the reads through filtering, dereplicating, removing chimeras, and constructing an amplicon sequence variant (ASV) abundance table. ASVs were annotated using the Bayesian LCA (BLCA) method (29) and the NCBI database. ASVs with BLCA confidence scores below 70% were labeled "unknown." For sets 1 and 3, ASVs linked to the genus *Lysobacter* were excluded due to its recognized presence as a contaminant in the achromopeptidase enzyme used during the DNA extraction process; for set 2, ASVs linked to *Lysobacter* were excluded for comparability to sets 1 and 3. Additionally, ASVs with total counts below the sample number threshold were excluded (n=48/52/54 for sets 1, 3, and 2, respectively). Contaminant ASVs were identified using control samples and were determined via a scoring method that encompasses results from Decontam (30), nonparametric statistical testing (Kruskal-Wallis *p*<0.5), mean comparison (i.e., mean counts of samples were less than mean counts of extraction controls), and whether sample read counts were less than the 5x greater than extraction controls counts. Decontaminated ASV counts were used for downstream analysis.

#### Analysis

For each set, the downstream analyses included beta diversity as measured by Principal Coordinate Analysis (PCoA) of decontaminated ASV counts. Following taxonomic identification, histograms and heatmaps were constructed showing the 49 most relatively abundant genera plus the categories “other” and “unknown.” With the histograms, we included total read count for each genus. These analyses were performed in R (R 4.3.1). For the PCoA, the Bray-Curtis distance was calculated using the vegdist function in the vegan library version 2.6.6.1 (31) and the resulting Bray-Curtis distance matrix was input to the pcoa function in the ape library version 5.8 to perform PCoA. The first two axes of the PCoA output were graphed using the plotly library version 4.10.4. The Bray-Curtis distance matrix was used as input to the PERMANOVA function in the PERMANOVA library version 0.2.0 to calculate the *p*-value. The relative abundance graphs were created by aggregating ASVs at the genus level and applying the transform function from the microbiome library (version 1.26.0) to transform the sample counts and visualizing the relative abundance using the plotly library (version 4.10.4). The heatmap analysis was graphed using the Heatmap function in the ComplexHeatmap library version 2.20.0 (32). To directly compare expressed urine samples from all three sets, the decontaminated ASV count tables from each set were combined into one database. Suprapubic aspirate samples were excluded. To directly compare paired suprapubic aspirates and expressed urine from the mice of set 3, the decontaminated ASV count tables were combined into one database. For both analyses, beta diversity, heatmap, and taxa histograms were performed as described above. For biplot analysis, genus level PCA was performed with the prcomp function (stats library v. 4.4.1) and visualized as a 3D biplot using factoextra (v.1.0.7).

## Results

We first determined a minimal urine volume for mouse urobiome studies, comparing 16S rRNA gene sequencing results across a range of urine volumes aliquoted from the same pooled urine from 4 different female mice. We determined that 500 μL was the smallest volume sufficient to extract and sequence enough DNA to adequately assess the urobiome while accounting for urine collection feasibility **(Supporting Fig. 1).**

**Supporting Figure 1. Determination of the smallest volume sufficient to extract and sequence enough DNA to adequately assess the urobiome while accounting for urine collection feasibility.** Histogram showing the 49 most relatively abundant genera detected in the extraction controls and expressed urine from 5 groups of mice. The category “Other” is an amalgamation of all other genera; the category “Unknown” represent all sequences that could not be identified. For each group of mice, different urine volumes were tested (125, 250, 500, 1000 μL). Note that the yellow line, which depicts total normalized arbitrary units (i.e., total reads per sample), correlates with sample volume, with 125μL producing numbers of reads apporximately the same as the extraction controls, while 1000 μL produced the largest number of reads. 500 μL were chosen for subsequent study as producing sufficiently high read numbers while balancing the feasiblity of obtaining urine from mouse bladders.

To determine whether the bioinformatically decontaminated data from each set of mice differed from their respective extraction controls, we performed PCoA using Bray-Curtis distances. For example, see **Supporting Fig. 2** for the analysis of the expressed urine samples of the set 1 mice. Although the samples differed significantly from the extraction controls (PERMANOVA, p = 0.003), some samples appeared similar to the controls. However, a histogram of aggregated ASVs at the genus level and total read counts revealed substantial differences in terms of composition with total read counts of samples 10- to 100-fold greater than those of the extraction controls (**Supporting Fig. 3**). Based on these results, we decided to include all samples in the downstream analyses.

**Supporting Figure 2. Example beta (between sample) diversity analysis used to determine whether expressed urine samples from set 1 (blue) differ from extraction controls (red).** Several samples appear to be quite similar to the extraction controls.

**Supporting Figure 3. Example histogram comparing the compositions of samples and extraction controls from set 1.** The histogram shows the 49 most relatively abundant genera plus plus the categories “other” and “unknown” detected in the samples and extraction controls from Supplemental Figure 2. Left Y-axis, relative abundance; right Y-axis, log_10_ of total units (i.e., reads). Note that the compositions of the samples differ from those of the extraction controls and that the samples contain at least 10-fold more reads than the controls. On the basis of this analysis, we decided to keep all samples for downstream analyses.

As a first step towards determining if an indigenous murine urobiome exists, we first determined whether the expressed samples from the three sets of mice contained bacteria (**Fig. 1**, **Fig. 2**). **Figure 1** is a set of three histograms (one for each set of mice), showing the 49 most relatively abundant genera across all three sets of mice, plus the categories “other” and “unknown.” **Figure 2** is a heatmap that presents the expressed urine samples from all three sets of mice. Taken together, these two figures show that the expressed urines contained DNA evidence of bacteria. The most prevalent and predominant genus in all three sets was *Staphylococcus.* Other less prevalent but predominant genera included *Enterococcus* (all sets) and *Mammaliicoccus* (sets 1 and 3).

**Figure 1.**
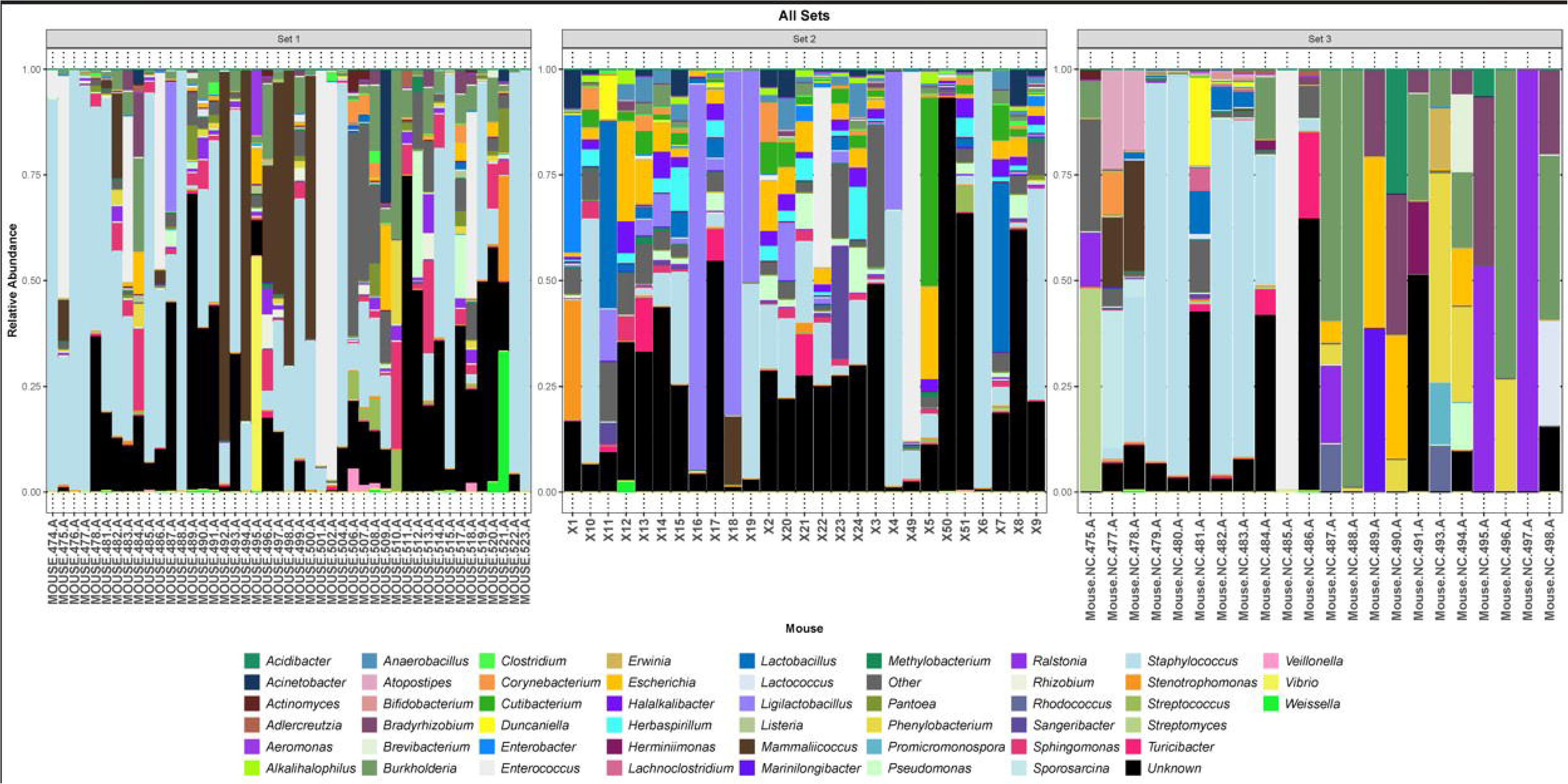
Three histograms showing the 49 most relatively abundant genera detected in each of the 3 sets of mice. The category “Other” is an amalgamation of all other genera; the category “Unknown” represent all sequences that could not be identified.

**Figure 2.**
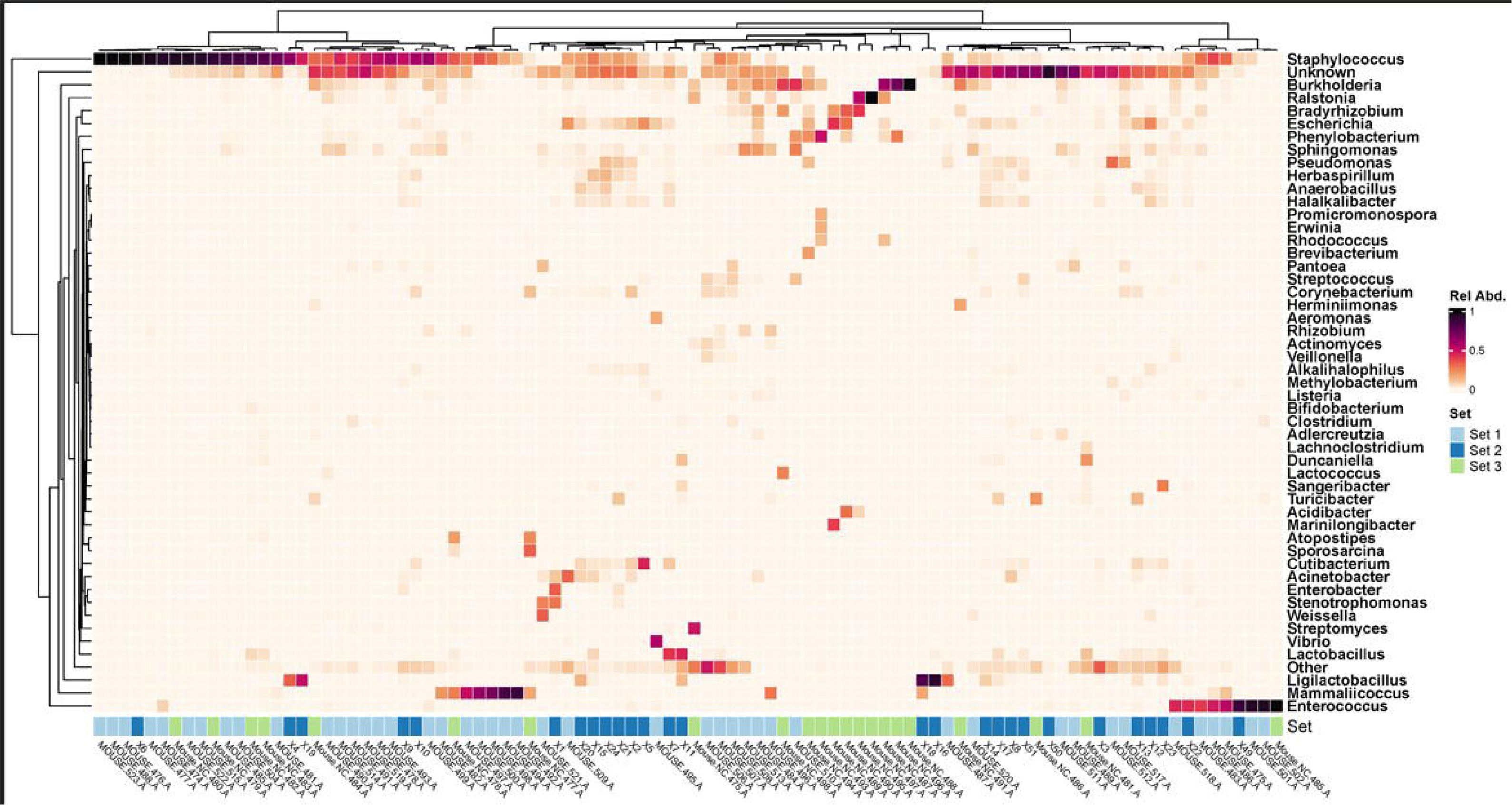
Heatmap showing the 49 most relatively abundant genera detected in the 3 sets of mice. The category “Other” is an amalgamation of all other genera; the category “Unknown” represent all sequences that could not be identified. Set 1, light blue; set 2, dark blue; set 3, green.

To compare the compositions of the expressed samples, we performed a PCoA. The three sets differed significantly whether tested together (**Fig. 3**, PERMANOVA, p = 0.001) or pairwise (**Supporting Fig. 4A-C**, PERMANOVA, p = 0.001 for all comparisons). Thus, the young mice housed in 2 different facilities (10 weeks of age; sets 1 and 2) differed significantly as did the young mice (set 1) and aged mice (6 ½ months of age; set 3). However, there appeared to be substantial overlap between the three sets. Sets 1 and 2 (both which included samples from young mice) had considerable overlap (**Supporting Fig. 4A**), as did sets 1 and 3 (young and aged mice) (**Supporting Fig. 4C**). Even sets 2 and 3 had some overlap (**Supporting Fig. 4C)**.

**Figure 3.**
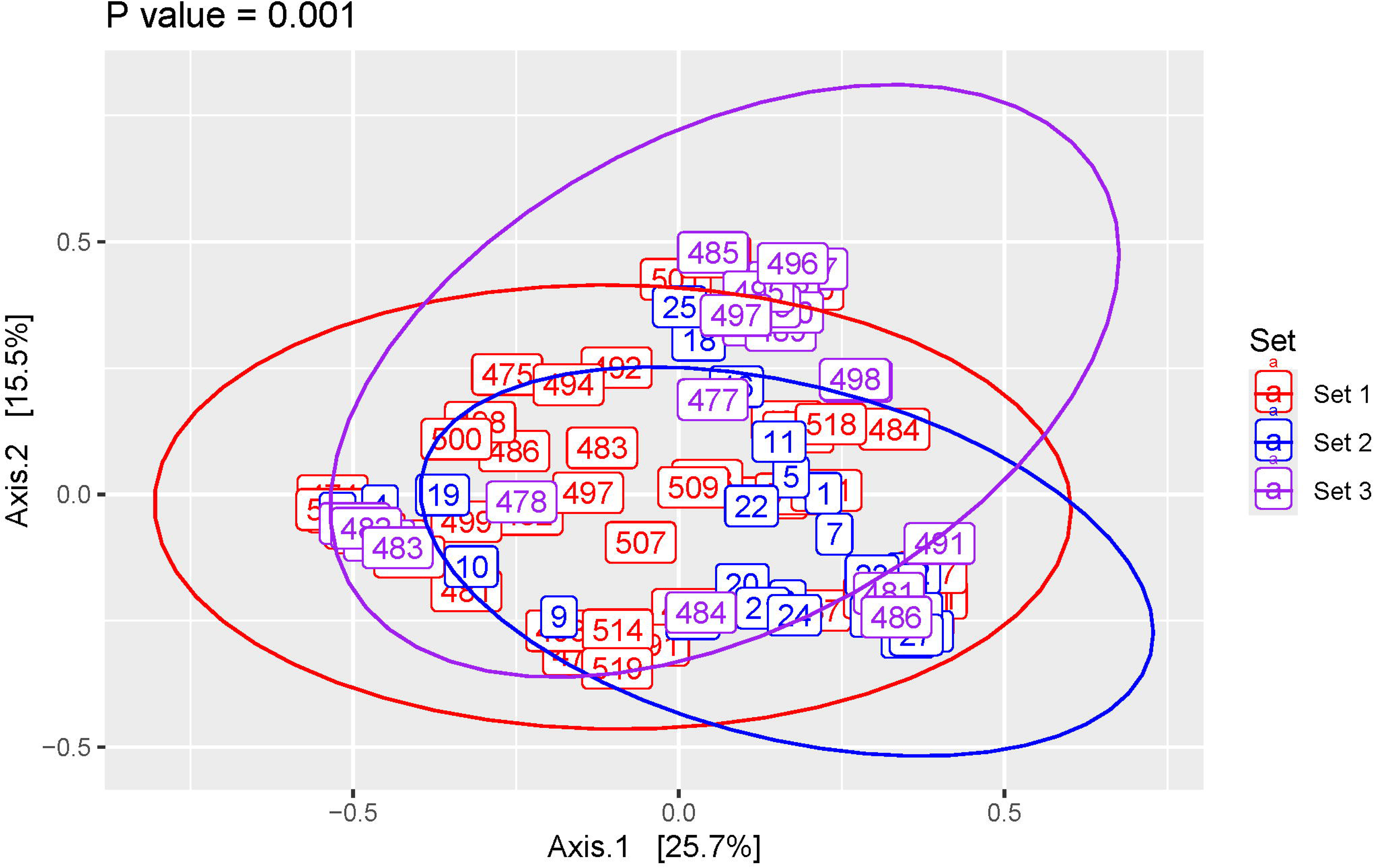
Beta (between participant) diversity analysis comparing all 3 sets of mice. Principal coordinate analysis of Bray-Curtis distances with P-value calculated using PERMANOVA.

To identify the genera most responsible for distinguishing each set from the others, we performed a biplot analysis, which identified them as *Staphylococcus, Mammaliicoccus, Ligilactobacillus, Enterococcus, Ralstonia, Burkholderia, Phenylobacterium, Escherichia,* and *Bradyrhizobium* plus the categories “other” and “unknown” (**Fig. 4**).

**Figure 4.**
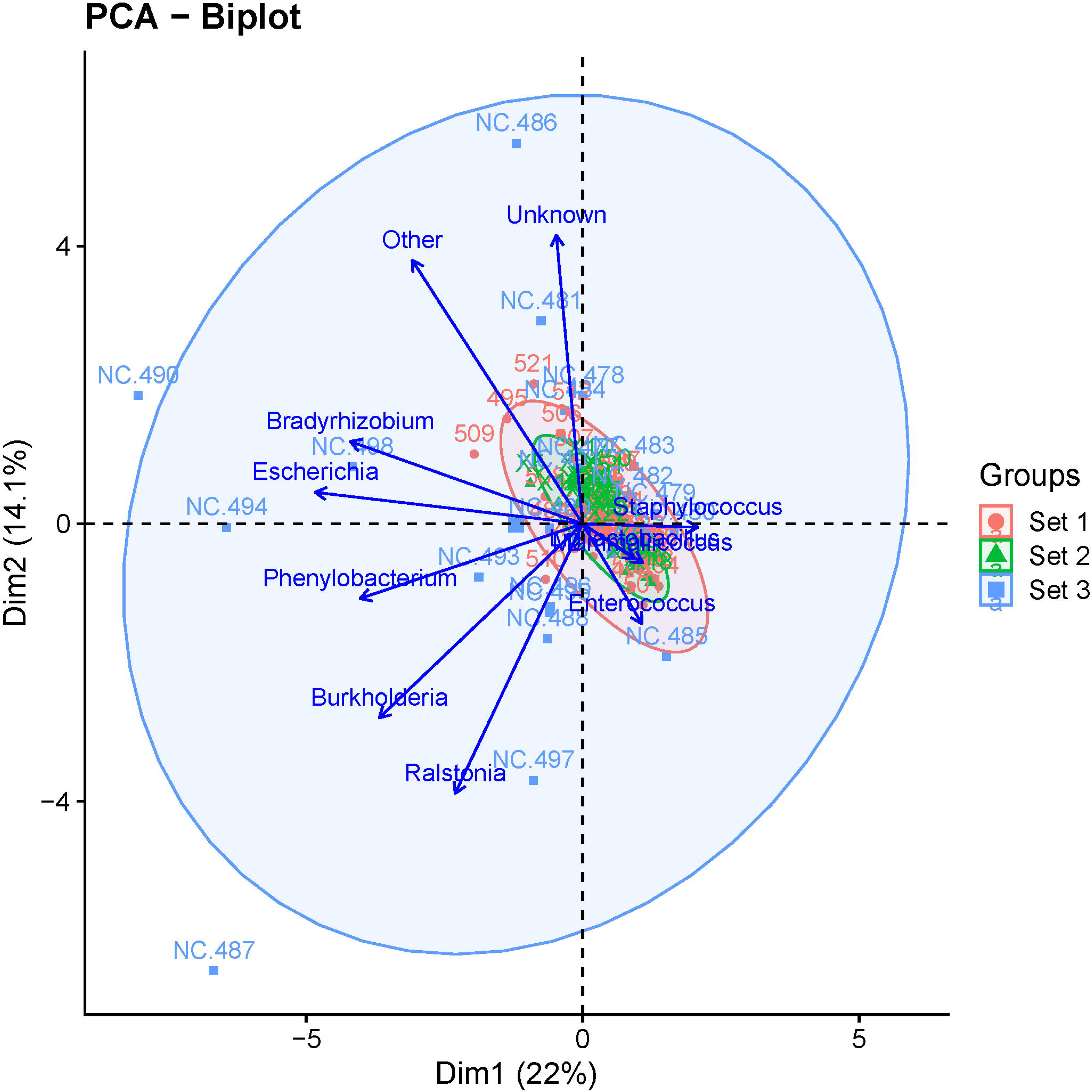
Biplot analysis identifying the taxa that differentiate the 3 sets of mice from each other.

**Supporting Figure 4. Beta (between participant) diversity analysis doing a pairwise comparison of all 3 sets of mice.** Principal coordinate analysis of Bray-Curtis distances with P-value calculated using PERMANOVA. (A) Set 1 versus set 2, (B) set 1 versus set 3, and (C) set 2 versus set 3.

Because some of these genera are considered to be environmental species, we hypothesized that expressed urine contained post-urethral microbes and thus compared the expressed urine samples of set 3 with their paired aspirated samples. First, we determined that the decontaminated aspirated samples differed significantly from the six extraction controls (**Supporting Fig. 5A**, PERMANOVA, p=0.001). One sample (483) seemed to resemble some of the controls, but when a genus level histogram was constructed, it differed sufficiently to be included downstream analyses (**Supporting Fig. 5B**). Next, we compared the two sample types, which differed significantly as determined by PCoA (**Fig. 5A**, PERMANOVA, p=0.001). This also was true when comparing the relative abundance of taxa between each pair of samples (**Fig. 5B**). The differences in bacterial taxa between aspirated and expressed samples are also illustrated in the heatmap (**Fig. 5C**). While the samples with the highest relative abundance of potential post-urethral microbes (e.g., *Staphylococcus, Ralstonia, Burkholderia)* are from expressed samples, some aspirated samples also contain these microbes, although at lower relative abundance compared to expressed samples.

**Figure 5.**
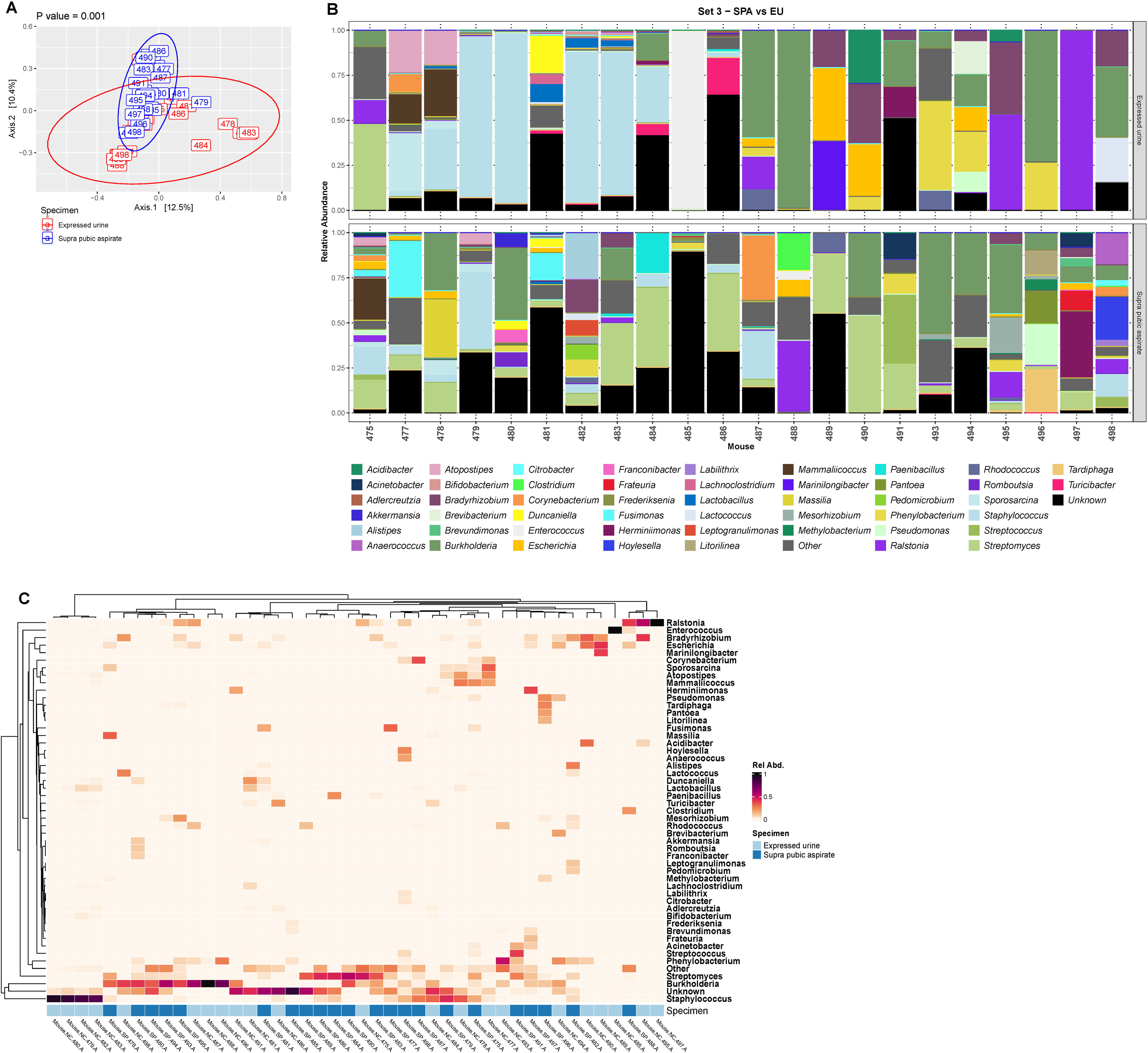
Comparison of urobiome composition in expressed urine versus paired suprapubic aspirate from the mice in set 3. (A) Principal coordinate analysis of Bray-Curtis distances with P-value calculated using PERMANOVA. (B) Histograms showing the 49 most relatively abundant genera detected in the paired expressed urine (top) and suprapubic aspirate (bottom) for each mouse in set 3. The category “Other” is an amalgamation of all other genera; the category “Unknown” represent all sequences that could not be identified. (C) Heatmap showing the 49 most relatively abundant genera detected in the paired expressed urine and suprapubic aspirate for each mouse in set 3. The category “Other” is an amalgamation of all other genera; the category “Unknown” represent all sequences that could not be identified. Light blue, expressed urine; dark blue, suprapubic aspirate.

**Supporting Figure 5. Comparison of suprapubic aspirate samples to extraction controls for set 3.** (A) Principal coordinate analysis of Bray-Curtis distances with P-value calculated using PERMANOVA. Note that one sample (483) appears to resemble the extraction controls. (B) Histogram showing the 49 most relatively abundant genera plus the categories “other” and “unknown” detected in the samples and extraction controls from panel A. Left Y-axis, relative abundance; right Y-axis, log_10_ of total units (i.e., reads). Note that the composition of sample 483 differed from those of the extraction controls. On the basis of this analysis, we decided to keep all samples for downstream analyses.

## Discussion

Here, we provide the first comprehensive report of the female mouse urobiome. We document similarity between the urobiome from the same strain of mice housed at different institutions, and an effect of age on the urobiome. Finally, we used suprapubic aspiration to demonstrate that there is a mouse urobiome that persists after removal of suspected contaminants, and that samples collected via aspiration significantly differ from those collected by manual expression.

An important aspect of this work is the data necessary to support the design of future research involving the mouse urobiome. First, we demonstrated that the urobiome is relatively consistent between mice housed in different sites. The mice in sets 1 and 2 were located in different institutions and had different facilities perform both the DNA extraction and sequencing. Despite these differences and significant differences based on PCoA analysis, there was substantial overlap between these 2 groups, suggesting some similarity between the sets. Second, we determined that 0.5 mL is an ideal urine volume for mouse urobiome studies. While this may not always be feasible and requires pooling of urine collected at different time points, this volume yielded sufficient sequencing depth. Third, we showed that aged mice before the onset of reproductive transition (∼6 ½ months) have urobiomes that differ significantly from their younger adult selves (10 weeks) but with substantial similarities. Finally, we showed that urine collected by manual expression has a high proportion of microbes from post-urethral sources and that collection of urine by a method that minimizes the risk of contamination, including aspiration, will reduce the inclusion of these microbes in the urine sample. Thus, when feasible, urine for urobiome studies from mice should be collected using a method that minimizes the potential for contamination, such as suprapubic aspiration.

Our work shows a small effect of age on the mouse urobiome. This is consistent with results from adult women, where the urobiomes of younger continent adult females were more likely to be predominated by members of the genus *Gardnerella*, whereas older continent adult females were more likely to be predominated by *Escherichia* (33). The authors hypothesized that the differences in the urobiome that corresponded to age were attributable to hormonal changes associated with aging. The mice in our work, while older, had not yet reached the age of reproductive transition, which correlates to menopause in women. This may partially explain the observed differences, but still there is a substantial overlap between our aged and non-aged mice. Further work that involves mice past the age of reproduction will provide further insights on the effect of age and hormonal status on the mouse urobiome.

There is limited literature with which to compare our results. However, there are a few works that document the rat urobiome. For example, Liu et al. (16) examined the effects of antibiotics on the urobiome of female rats. They collected urine by urethral catheterization and used 16S rRNA gene sequencing to describe the urobiome. Their control animals, which did not have any interventions, had a urobiome. While there are several genera in common between the urobiome of their control animals and our mice, it is notable that their most abundant genus, *Bacillus*, was not within the 51 most abundant genera that we identified in mice. Chen et al. also examined the rat urobiome but used autoclaved metabolic cages for urine collection (17). Their genus level microbiome results from their female control rats have several genera in common with what we report in mice, including *Escherichia, Streptococcus,* and *Acinetobacter.* Thus, while not directly comparable given the differences in species, these reports that document the urobiome in rats lend credence to our data documenting the urobiome in mice.

One of the most relevant implications of our work, as well as our documentation of the presence of the host urobiome, is related to preclinical *in vivo* UTI work. There is a vast body of preclinical studies using the mouse model of UTI that have yielded critical information about the pathobiology of uropathogenic *Escherichia coli* and host immune responses related to UTI (10, 24, 34). However, a notable aspect of this model is that mice require a large number of bacteria, usually 10^7^ colony forming units of *E. coli,* to induce more than transient bacteriuria. Furthermore, not all mice that have that large number of bacteria instilled into their bladders develop bacteriuria. One previously overlooked factor into the heterogeneity of UTIs in mice is the native urobiome and its role in colonization resistance. The concept of colonization resistance has been described in the context of probiotics in the gut microbiome in which features of the gut microbiome can predict subsequent colonization by exogenous bacteria. Colonization resistance is built on the concept that a microbiome is an ecologically stable community of bacteria (35). Disrupting an ecologically stable community requires alteration of the host microbiome such as through exposure to antimicrobials, alteration of the host to affect availability of resources, or provision of a large influx of exogenous microbes that subsequently outcompetes the native community, such as occurs in the UTI model. Considering the role of the native urobiome in UTI susceptibility is critical to development of more translatable models (22, 23).

Limitations of this work include a small sample size of mice in each group. Furthermore, we only included female C57BL/6 mice, and our results cannot be generalized to either male mice or female mice of other strains. It is likely that both sex- and strain-based differences exist within the urobiome that should be described in future work. Also, we used short-read (V4) 16S rRNA gene sequencing, which typically yields only confidently identifies taxonomy at the family or genus levels. Future work should include long read 16S rRNA gene sequencing for confident species level identification. Despite these limitations, our work does have several strengths. We obtained data from two different sets of mice in two different facilities, used two different sequencing methods, but analyzed all the sequence data simultaneously with a single analytic approach. We compared expressed versus aspirated urine to describe the effects of post-urethral microbes on expressed urobiome samples. We used an improved DNA isolation method that uses an extra enzyme (lysostaphin) with longer incubation times to increase cell wall breakage. Finally, in the analytic step, we used a multi-step decontamination procedure.

## Conclusions

There is a native mouse urobiome that is largely stable in the same strain of mice. There is a small effect of age in female mice who are capable of reproduction. Finally, use of suprapubic aspiration, or a similar technique, to bypass post-urethral microbes, is critical for accurately assessing the mouse urobiome.

## Author contributions

AJW and CSF designed and implemented the original study protocol and obtained funding. DB, SN, OL, and BS did the mouse work. MK processed the samples for sequencing. SS analyzed the data. MK and SS constructed the figures. AJW, MK, and CSF interpreted data. AJW, CSF, MK, RBM, and SS wrote the manuscript. All authors reviewed and approved the manuscript.

## Disclosure statement from authors

AJW discloses membership on the Scientific Advisory Boards of Urobiome Therapeutics, Pathnostics, and Cerillo. He also discloses funding from the Craig H. Neilsen Foundation, Pathnostics, and an anonymous donor. The rest of the authors have no relevant disclosures.

## Data availability

Raw sequencing reads are publicly available at the NCBI SRA database (BioProject ID: PRJNA1229522).

## Acknowledgements

We gratefully acknowledge Melline Fontes Noronha for help with bioinformatics.

## References

1. Moreland RB, Brubaker L, Tinawi L, Wolfe AJ. Rapid and accurate testing for urinary tract infection: new clothes for the emperor. Clin Microbiol Rev. 2025;38(1):e0012924.

2. Simoni A, Schwartz L, Junquera GY, Ching CB, Spencer JD. Current and emerging strategies to curb antibiotic-resistant urinary tract infections. Nat Rev Urol. 2024;21:707–22.

3. MacDonald RA, Levitin H, Mallory GK, Kass EH. Relation between pyelonephritis and bacterial counts in the urine. N Engl J Med. 1957;256(20):915–22.

4. JawetzE. Urinary tract infections; problems in medical management. California Med. 1953;79(2):99–102.

5. Timm MR Russell SK, Hultgren SJ. Urinary tract infections: pathogenesis, host susceptibility and emerging therapeutics. Nat Rev Microbiol. 2025;23(2):72–86.

6. Brubaker L Chai T, Horsley H, Khasriya R, Moreland RB and Wolfe AJ. Tarnished gold—the “standard” urine culture: reassessing the characteristics of a criterion standard for detecting urinary microbes. Front Urol. 2023;3:1206046.

7. Wolfe AJ, Brubaker L. Discovering the Urinary Microbiome. American Scientist. 2024;112(3):168–75.

8. Price TK, Dune T, Hilt EE, Thomas-White KJ, Kliethermes S, Brincat C, Brubaker L, Wolfe AJ, Mueller ER, Schreckenberger PC. The Clinical Urine Culture: Enhanced Techniques Improve Detection of Clinically Relevant Microorganisms. J Clin Microbiol. 2016;54(5):1216–22.

9. Du J, Khemmani M, Halverson T, Ene A, Limeira R, Tinawi L, Hochstedler-Kramer BR, Noronha MF, Putonti C, Wolfe AJ. Cataloging the phylogenetic diversity of human bladder bacterial isolates. Genome Biol. 2024;25(1):75.

10. Murray BO, Flores C, Williams C, Flusberg DA, Marr EE, Kwiatkowska KM, Charest JL, Isenberg BC, Rohn JL. Recurrent Urinary Tract Infection: A Mystery in Search of Better Model Systems. Front Cell Infect Microbiol. 2021;11:691210.

11. Burton EN, Cohn LA, Reinero CN, Rindt H, Moore SG, Ericsson AC. Characterization of the urinary microbiome in healthy dogs. PLoS One. 2017;12(5):e0177783.

12. Coffey EL, Gomez AM, Burton EN, Granick JL, Lulich JP, Furrow E. Characterization of the urogenital microbiome in Miniature Schnauzers with and without calcium oxalate urolithiasis. J Vet Intern Med. 2022;36(4):1341–52.

13. Coffey EL, Gomez AM, Ericsson AC, Burton EN, Granick JL, Lulich JP, Furrow E. The impact of urine collection method on canine urinary microbiota detection: a cross-sectional study. BMC Microbiol. 2023;23(1):101.

14. Coffey EL, Becker ZW, Gomez AM, Ericsson AC, Churchill JA, Burton EN, Granick JL, Lulich JP, Furrow E. Dietary Features Are Associated with Differences in the Urinary Microbiome in Clinically Healthy Adult Dogs. Vet Sci. 2024;11(7):286.

15. Joubran P, Roux FA, Serino M, Deschamps JY. Gut and Urinary Microbiota in Cats with Kidney Stones. Microorganisms. 2024;12(6):1098.

16. Liu F HL, Sheng J, Sun Y, Xia Q, Tang Y, Jiang P, Wei S, Hu J, Lin H, Xu Z, Guo W, Gu Y, Feng N. Can antibiotics for enteritis or for urinary tract infection disrupt the urinary microbiota in rats? Front Cell Infect Microbiol. 2023;28(13):1169909.

17. Chen X, Cheng Y, Tian X, Li J, Ying X, Zhao Q, Wang M, Liu Y, Qiu Y, Yan X, Ren X. Urinary microbiota and metabolic signatures associated with inorganic arsenic-induced early bladder lesions. Ecotoxicol Environ Saf. 2023;259:115010.

18. Karstens L Asquith M, Caruso V, Rosenbaum JT, Fair DA, Braun J, Gregory WT, Nardos R, McWeeney SK. Community profiling of the urinary microbiota: considerations for low-biomass samples. Nat Rev Urol. 2018;15(12):735–49.

19. Agudelo J, Miller AW. Impact of study design, contamination, and data characteristics on results and interpretation of microbiome studies. mSystems. 2025; Online ahead of print.

20. Bleich A FJ. The Mammalian Microbiome and Its Importance in Laboratory Animal Research. ILAR J. 2015;56(2):153–8.

21. Jayalath S, Magana-Arachchi D. Dysbiosis of the Human Urinary Microbiome and its Association to Diseases Affecting the Urinary System Indian J Microbiol. 2022;62(2):133–66.

22. Pastuszka A, Tobor S, Łoniewski I, Wierzbicka-Woś A, Sielatycka K, Styburski D, Cembrowska-Lech D, Koszutski T, Kurowicz M, Korlacka K, Podkówka A, Lemiński A, Brodkiewicz A, Hyla-Klekot L, Skonieczna-Żydecka K. Rewriting the urinary tract paradigm: the urobiome as a gatekeeper of host defense. Mol Biol Rep. 2025;52(1):497.

23. Reasoner SA, Francis J, Hadjifrangiskou M. The urinary microbiome: the next frontier of bacterial ecology. J Bacteriol. 2025;:e0010525(:e0010525): Online ahead of print.

24. Carey AJ, Tan CK, Ipe DS, Sullivan MJ, Cripps AW, Schembri MA, Ulett GC. Urinary tract infection of mice to model human disease: Practicalities, implications and limitations. Crit Rev Microbiol. 2016;42(5):780–99.

25. Anderson M, Bollinger D, Hagler A, Hartwell H, Rivers B, Ward K, Steck TR. Viable but nonculturable bacteria are present in mouse and human urine specimens. J Clin Microbiol. 2004;42(2):753–8.

26. Roje B, Elek A, Palada V, Bom J, Iljazović A, Šimić A, Sušak L, Vilović K, Strowig T, Vlahoviček K, Terzić J. Microbiota Alters Urinary Bladder Weight and Gene Expression. Microorganisms. 2020;8(3):421.

27. Martin M. CUTADAPT removes adapter sequences from high-throughput sequencing reads. EMBNet J. 2011;17(1):10–2.

28. Callahan BJ, McMurdie PJ, Rosen MJ, Han AW, Johnson AJ, Holmes SP. DADA2: High-resolution sample inference from Illumina amplicon data. Nat Methods. 2016;13(7):581–3.

29. Gao X, Lin H, Revanna K, Dong Q. A Bayesian taxonomic classification method for 16S rRNA gene sequences with improved species-level accuracy. BMC Bioinformatics. 2017;18(1):247.

30. Davis NM, Proctor DM, Holmes SP, Relman DA, Callahan BJ. Simple statistical identification and removal of contaminant sequences in marker-gene and metagenomics data. Microbiome. 2018;6(1):226.

31. Oksanen J, Simpson GL, Blanchet FG, Kindt R, Legendre P, Minchin PR, O’Hara RB, Solymos P, Stevens MHM, Szoecs E, Wagner H, Barbour M, Bedward M, Bolker B, Borcard D, Borman T, Carvalho G, Chirico M, De Caceres M, Durand S, Evangelista HBA, FitzJohn R, Friendly M, Furneaux B, Hannigan G, Hill MO, Lahti L, Martino C, McGlinn D, Ouellette MH, Cunha ER, Smith T, Stier A, Ter Braak JF, Weedon J. vegan: Community Ecology Package. R package version 2.2–1 2015 [Available from: http://CRAN.R-project.org/package=vegan.

32. Gu Z, Miller S, Schlesner M. Complex heatmaps reveal patterns and correlations in multidimensional genomic data. Bioinformatics. 2016;32(18):2847–9.

33. Price TK, Hilt EE, Thomas-White K, Mueller ER, Wolfe AJ, Brubaker L. The urobiome of continent adult women: a cross-sectional study. BJOG. 2020;127(2):193–201.

34. Chieng CCY, Kong Q, Liou NSY, Khasriya R, Horsley H. The clinical implications of bacterial pathogenesis and mucosal immunity in chronic urinary tract infection. Mucosal Immunol. 2023;16(1):61–71.

35. Wu G, Zhao N, Zhang C, Lam YY, Zhao L. Guild-based analysis for understanding gut microbiome in human health and diseases. Genome Med. 2021;13(1):22.

