## Supporting Information Fig 1-5 for "Evidence for an indigenous female mouse urobiome"

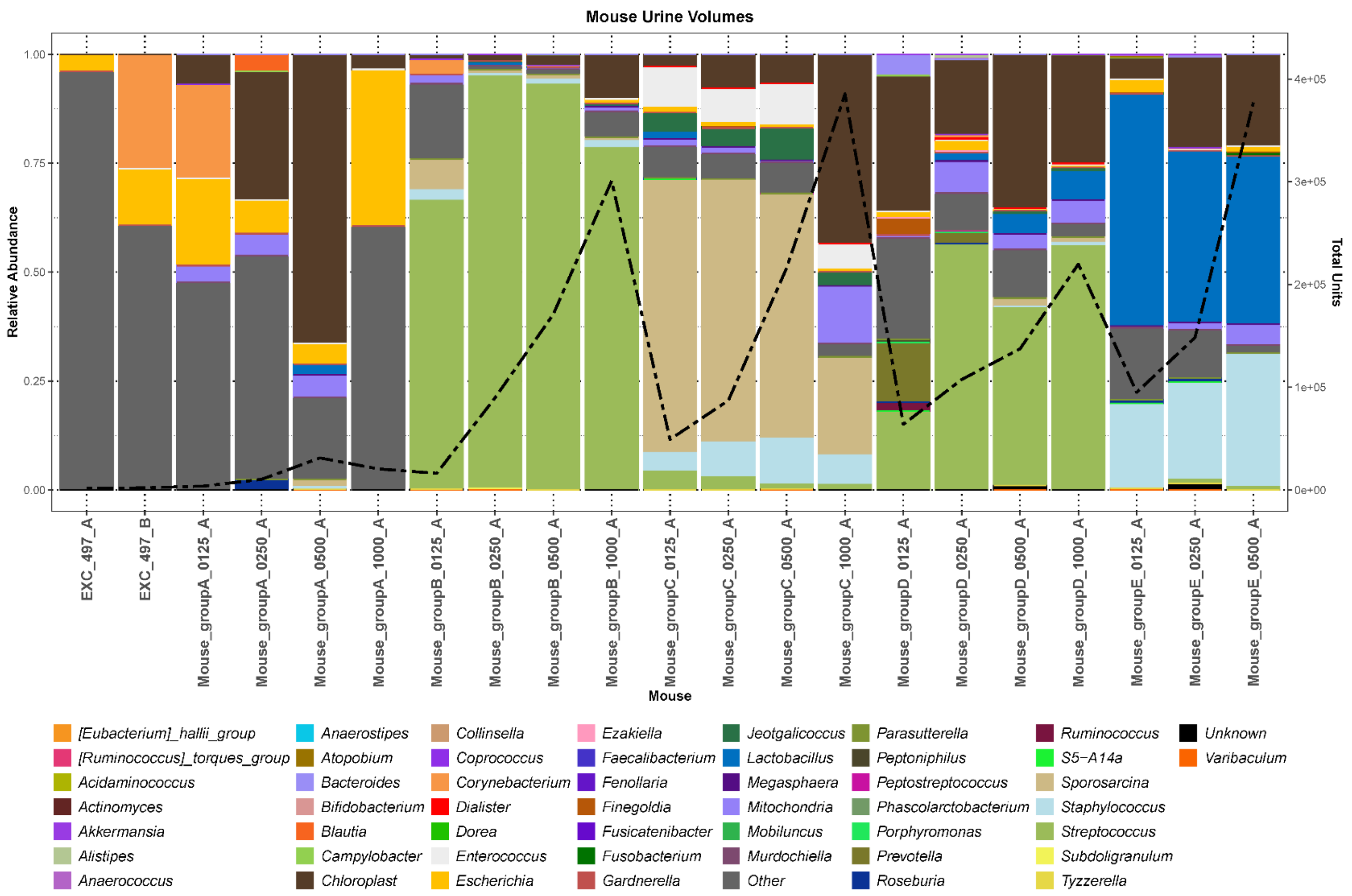


**Supporting Figure 1.** Determination of the smallest volume sufficient to extract and sequence enough DNA to adequately assess the urobiome while accounting for urine collection feasibility. Histogram showing the 49 most relatively abundant genera detected in the extraction controls and expressed urine from 5 groups of mice. The category "Other" is an amalgamation of all other genera; the category "Unknown" represent all sequences that could not be identified. For each group of mice, different urine volumes were tested (125, 250, 500, 1000 μL). Note that the yellow line, which depicts total normalized arbitrary units (i.e., total reads per sample), correlates with sample volume, with 125μL producing numbers of reads approximately the same as the extraction controls, while 1000 μL produced the largest number of reads. 500 μL were chosen for subsequent study as producing sufficiently high read numbers while balancing the feasibility of obtaining urine from mouse bladders.

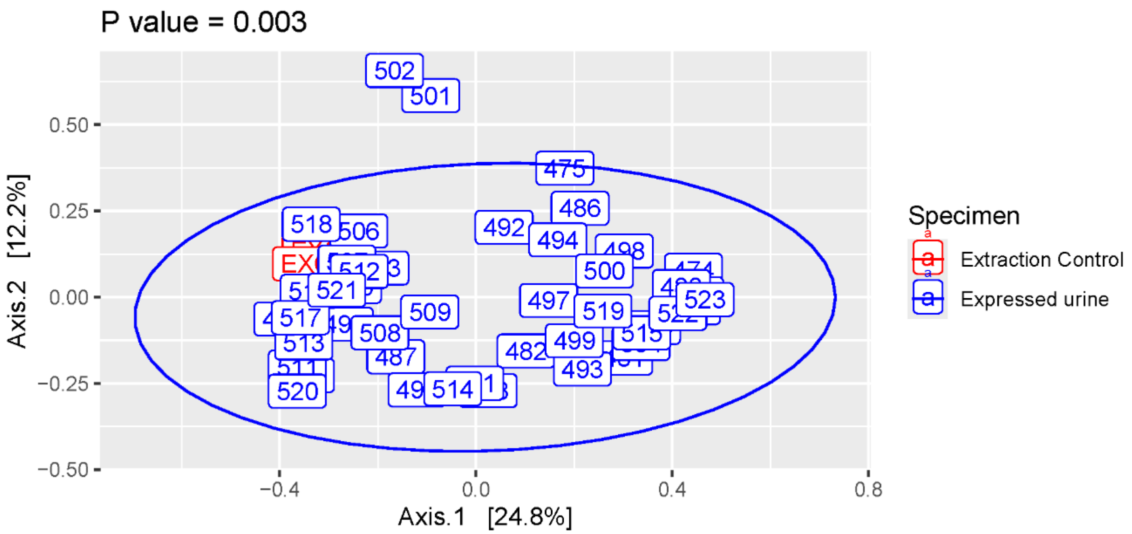


**Supporting Figure 2**. Example beta (between sample) diversity analysis used to determine whether expressed urine samples from set 1 (blue) differ from extraction controls (red). Several samples appear to be quite similar to the extraction controls.

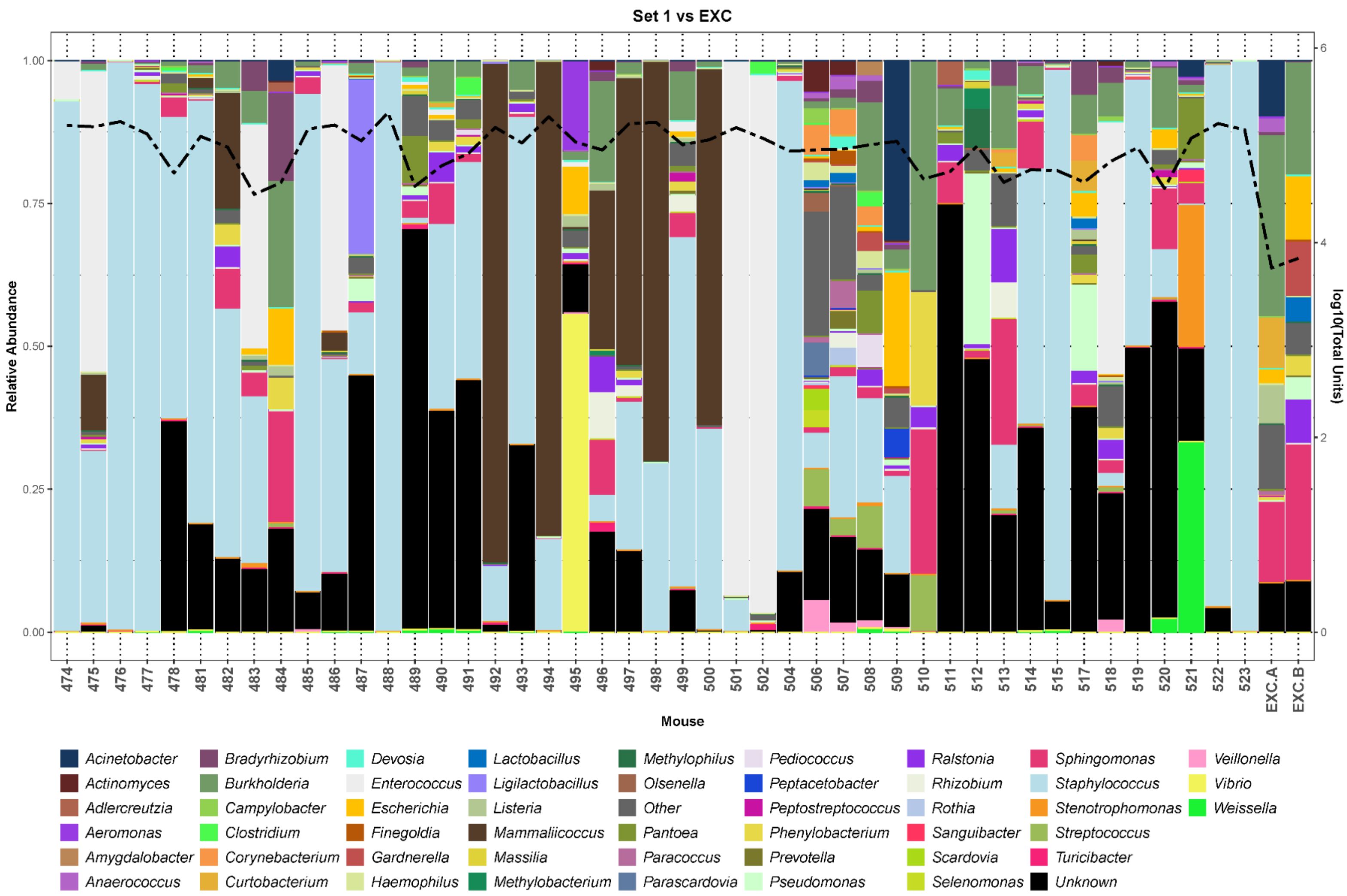


**Supporting Figure 3**. Example histogram comparing the compositions of samples and extraction controls from set 1. The histogram shows the 49 most relatively abundant genera plus plus the categories “other” and “unknown” detected in the samples and extraction controls from Supplemental Figure 2. Left Y-axis, relative abundance; right Y-axis, log10 of total units (i.e., reads). Note that the compositions of the samples differ from those of the extraction controls and that the samples contain at least 10-fold more reads than the controls. On the basis of this analysis, we decided to keep all samples for downstream analyses.

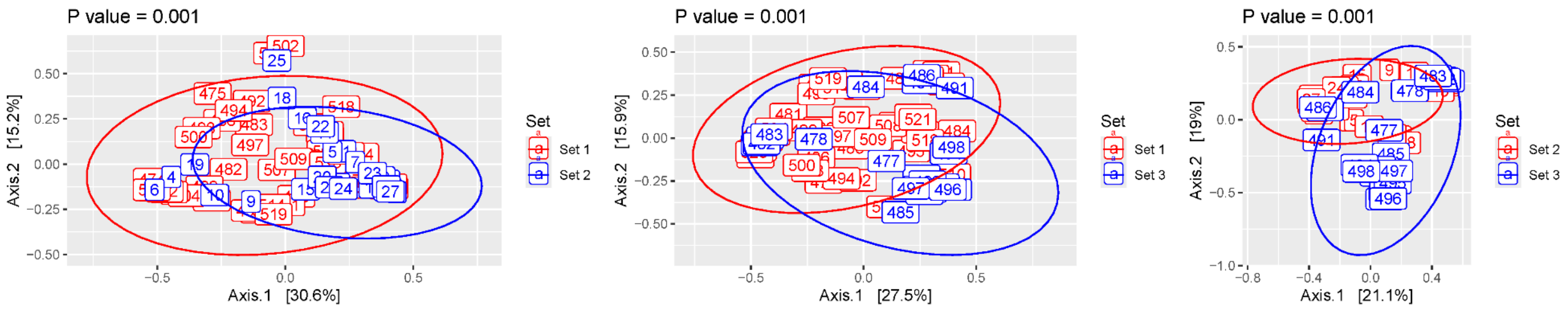


**Supporting Figure 4**. Beta (between participant) diversity analysis doing a pairwise comparison of all 3 sets of mice. Principal coordinate analysis of Bray-Curtis distances with P-value calculated using PERMANOVA. (A) Set 1 versus set 2, (B) set 1 versus set 3, and (C) set 2 versus set 3.

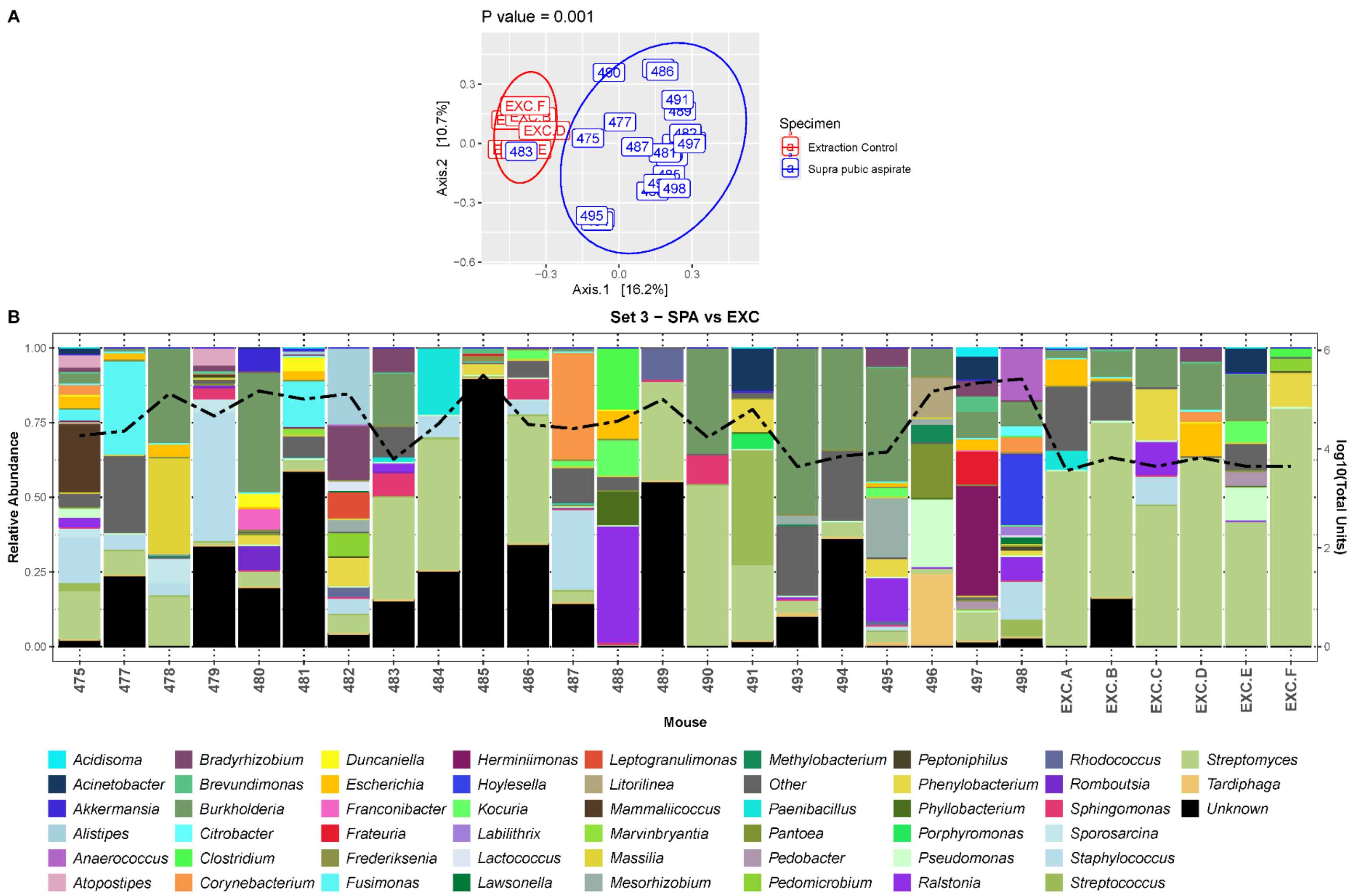


;

**Supporting Figure 5.** Comparison of suprapubic aspirate samples to extraction controls for set 3. (A) Principal coordinate analysis of Bray-Curtis distances with P-value calculated using PERMANOVA. Note that one sample (483) appears to resemble the extraction controls. (B) Histogram showing the 49 most relatively abundant genera plus the categories "other" and "unknown" detected in the samples and extraction controls from panel A. Left Y-axis, relative abundance; right Y-axis, log10 of total units (i.e., reads). Note that the composition of sample 483 differed from those of the extraction controls. On the basis of this analysis, we decided to keep all samples for downstream analyses.
